# Comparison of metabarcoding taxonomic markers to describe fungal communities in fermented foods

**DOI:** 10.1101/2023.01.13.523754

**Authors:** Olivier Rué, Monika Coton, Eric Dugat-Bony, Kate Howell, Françoise Irlinger, Jean-Luc Legras, Valentin Loux, Elisa Michel, Jérôme Mounier, Cécile Neuvéglise, Delphine Sicard

**Affiliations:** Université Paris-Saclay, INRAE, MaIAGE, 78350, Jouy-en-Josas, France; Université Paris-Saclay, INRAE, BioinfOmics, MIGALE bioinformatics facility, 78350, Jouy-en-Josas, France; Univ. Brest, INRAE, Laboratoire Universitaire de Biodiversité et Ecologie Microbienne, F-29280 Plouzané, France; Université Paris Saclay, INRAE, AgroParisTech, UMR SayFood, 91120 Palaiseau, France; School of Agriculture and Food, Faculty of Veterinary & Agricultural Sciences, The University of Melbourne, Parkville, Victoria, Australia; SPO, Univ Montpellier, INRAE, Institut Agro, Montpellier, France

**Keywords:** fermented foods, metabarcoding, fungi, mock communities, barcode marker comparison

## Abstract

Next generation sequencing offers several ways to study microbial communities. For agri-food sciences, identifying species in diverse food ecosystems is key for both food sustainability and food security. The aim of this study was to compare metabarcoding pipelines and markers to determine fungal diversity in food ecosystems, from Illumina short reads. We built mock communities combining the most representative fungal species in fermented meat, cheese, wine and bread. Four barcodes (ITS1, ITS2, D1/D2 and RPB2) were tested for each mock and on real fermented products. We created a database, including all mock species sequences for each barcode to compensate for the lack of curated data in available databases. Four bioinformatics tools (DADA2, QIIME, FROGS and a combination of DADA2 and FROGS) were compared. Our results clearly showed that the combined DADA2 and FROGS tool gave the most accurate results. Most mock community species were not identified by the RPB2 barcode due to unsuccessful barcode amplification. When comparing the three rDNA markers, ITS markers performed better than D1D2, as they are better represented in public databases and have better specificity to distinguish species. Between ITS1 and ITS2, differences in the best marker were observed according to the studied ecosystem. While ITS2 is best suited to characterize cheese, wine and fermented meat communities, ITS1 performs better for sourdough bread communities. Our results also emphasized the need for a dedicated database and enriched fungal-specific public databases with novel barcode sequences for 118 major species in food ecosystems.

## Introduction

In the field of microbial ecology, amplicon-based metagenomic analysis (also named metabarcoding) is one of the most popular routes to describe microbial communities as it is a high throughput method with relatively low-cost nowadays. This approach relies on amplifying a phylogenetic biomarker from total community DNA purified from the samples to be characterized, followed by massive amplicon sequencing (Shokralla et al., 2012), usually with Illumina MiSeq technology (Caporaso et al., 2012). For bacteria, the SSU rRNA gene is commonly accepted as the most suitable biomarker for metabarcoding studies although different variable regions can be targeted (Zhang et al., 2018). For fungal communities, there is still no international consensus regarding the choice of the best phylogenetic biomarker for such an approach.

Ten years ago, the Fungal Barcoding Consortium recommended the use of the Internal Transcribed Spacer (ITS) region as the primary marker for fungal identifications due to superior species-level resolution compared to LSU and SSU rRNA genes, and higher amplification success compared to protein coding genes (Schoch et al., 2012). However, amplicon-based metagenetic analysis using MiSeq sequencing imposes more constraints than classical species identification. Most importantly, the typical amplicon size (∼500bp) will not provide complete ITS region sequences or full length rRNA or protein coding genes. Consequently, the most popular barcodes currently used for fungal community analysis by metabarcoding are ITS1, ITS2 and the LSU D1/D2 domain (R. Henrik Nilsson et al., 2019).

Amplicon-based metagenetic analyses using these markers has been largely facilitated over the past few years by the availability of a broad range of high-quality reference sequences in public databases from different sequencing initiatives of collection strains (Vu et al., 2016).

However, the multicopy nature of the rDNA operon negatively affects fungal community compositions detected in complex samples by metabarcoding (Lavrinienko et al., 2021). In addition, using the entire ITS region barcode leads to additional bias resulting in lower representation of species with longer amplicons in the datasets (Ihrmark et al., 2012). Nevertheless, using only ITS1 or ITS2, which produce shorter amplicons, was shown to represent the quantitative composition of the sample (Ihrmark et al., 2012). Several very promising single-copy marker genes were thus proposed to overcome these limitations, including the rpb2 gene, encoding for the second largest ribosomal polymerase II subunit (Větrovský et al., 2016). In addition to the above-mentioned specificity, the species-resolving power of rpb2 was found to be higher than rDNA genes and ITS regions. This marker was also found to be particularly suited to study basal fungal lineages. Yet, due to the lack of universality and lower specificity of RPB2 primers (when compared to others) as well as the lower numbers of rpb2 gene sequences in public databases, further applications to study fungal communities from food ecosystems may be limited.

Some authors evaluated the reliability of different markers (ITS1-2, D1/D2 LSU and SSU) to describe fungal diversity by amplicon-based metagenomic analysis (De Filippis et al., 2017). A mock community, composed of 19 strains representative of common fungal species, as well as environmental samples including soil, human saliva, human feces and grape must were used. Although all markers were able to correctly detect the different species in the mock community, the results suggested that there was an important quantification bias when using ITS1-2. This could be due to the high heterogeneity in marker length across fungal species. However, marker performance is likely to be highly influenced by fungal species composition of the sample (e.g., composed mainly of Basidiomycota versus Ascomycota) which, in turn, depends on the studied environment.

Numerous tools are available for fungal metabarcoding data analyses from Illumina sequencing technology (R. Henrik Nilsson et al., 2019) but many differences between pipelines exist (Anslan et al., 2018) which highlights the need to pay attention to the choice of the tool. Indeed, amplicon length is one crucial characteristic to consider. Using an *in silico* approach for fungal sequences, a recent showed that the length of the extracted ITS1 portions from UNITE ranged from 9 bp to 1181 bp, with an average length of 177 bp while the extracted ITS2 portions ranged from 14 bp to 730 bp, with an average of 182 bp (Yang et al., 2018). For D1/D2 and RPB2 amplicons, lengths are often above 600 bp; in this case, common strategies that merge paired-end reads are not suitable because Illumina sequencing, the most used technology in metabarcoding analyses, provides paired-end reads of maximum 2 × 300 bp.

Among bioinformatics solutions for fungal communities with short reads, some are dedicated to ITS such as PIPITS (Gweon et al., 2015) or DAnIEL (Loos et al., 2021) while others are more generic (Bernard et al., 2021; Bolyen et al., 2019; Callahan et al., 2016; Edgar, 2010; Escudié et al., 2018; Özkurt et al., 2022). However, one downside is that few of them can process short and long amplicons simultaneously. PIPITS and DAnIEL reject non-overlapping reads, so long amplicons will be excluded. QIIME2 and DADA2 require a choice to be made between merging reads or keeping only R1 reads. USEARCH recommends taking into account merged sequences and 5′ R1 reads of non-overlapping paired-end sequences. FROGS deals with mergeable reads and creates artificial sequences from non-mergeable reads, therefore all sequenced information is kept throughout the pipeline. It is worth mentioning that FROGS has shown better results than QIIME2, DADA2 and USEARCH on simulated data (Bernard et al., 2021).

Another important aspect in metabarcoding data analysis is the way to build representative biological sequences from reads. It can be under the form of Operational Taxonomic Units (OTUs), Amplicon Sequence Variants (ASVs) or zero-radius OTUs (ZOTUs) depending on tools. Numerous studies have compared the results from OTU-based and ASV-based approaches (Callahan et al., 2017). Using mock communities, ASV-based methods had higher sensitivity and detected bacterial strains present, sometimes at the expense of specificity (Caruso et al., 2019). However, a different study concluded that for broadscale (e.g., all bacteria or all fungi) α and β diversity analyses, ASV or OTU methods often provided similar ecological results (Glassman and Martiny, 2018). From a practical point of view, an important advantage of ASV-based approaches is the consistent labels with intrinsic biological meaning identified independently from a reference database. Thus, ASVs independently inferred from different studies and different samples (for the same targeted region) can be compared.

In this study, we compared the performance of four phylogenetic markers (ITS1, ITS2, D1/D2 LSU and RPB2) for metabarcoding analysis of complex fungal communities in different fermented foods, after assessing which bioinformatic strategy was most suitable for analyzing such datasets. To perform these analyses, we compared seven strategies based on commonly used tools for metabarcoding data (QIIME2, DADA2, USEARCH, FROGS) and a combination of DADA2 and FROGS (named DADA2_FROGS) by analyzing four separate mock communities’ representative of the fungal diversity found in meat sausage (fermented meat), cheese, grape must (wine) and sourdough (bread), for a total of 118 species, as well as 24 real fermented food samples.

## Methods

### Mock community sample preparation

#### Species diversity

For each fermented food type, representative species were selected based on an inventory of the most frequently described species in the literature. One strain (mainly available type-strains) was included for each selected species. The complete list of strains used for mock community design is available on Recherche Data Gouv platform (https://doi.org/10.57745/AZNJFE) and a summary is presented in Table 1.

**Table 1.** Number of Species, Genus and Family in each mock community

|  | Bread | Cheese | Meat | Wine |
| --- | --- | --- | --- | --- |
| Species | 27 | 25 | 40 | 60 |
| Genus | 15 | 16 | 14 | 37 |
| Family | 4 | 11 | 8 | 8 |

Figure 1 shows the species distribution at the genus level in the different mock community samples.

**Figure 1.**
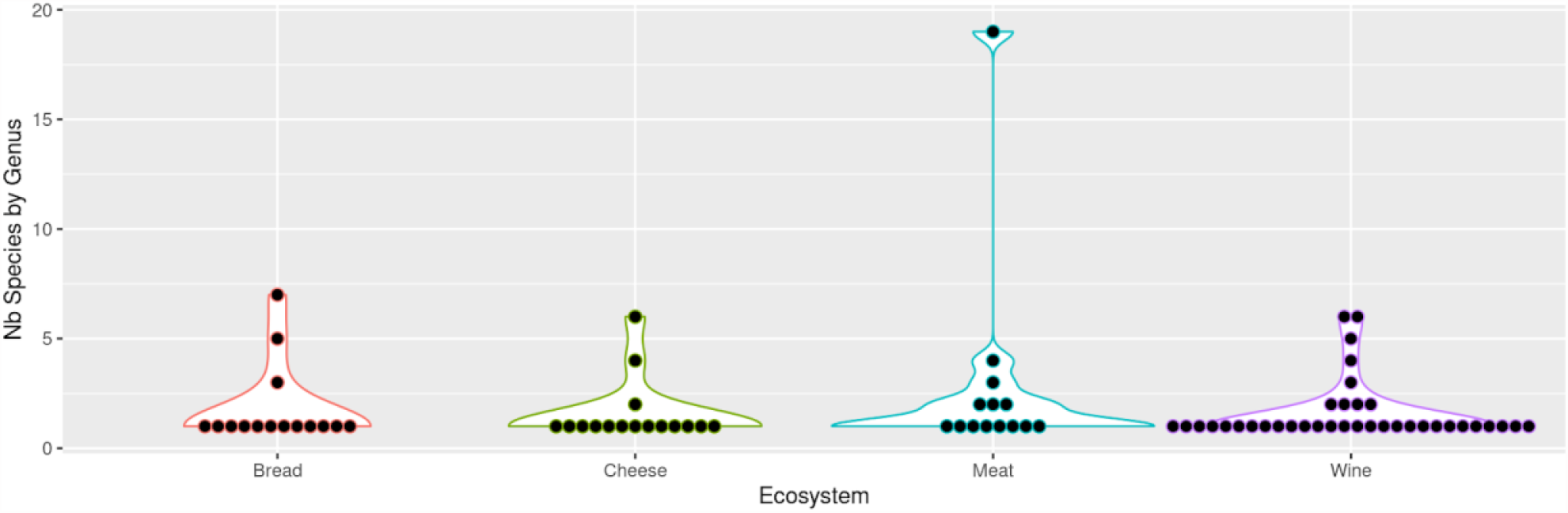
Genus and species diversity in mock community samples. Each point corresponds to a genus. The number of species per genus is indicated on the vertical axis. For example, in meat, one genus is represented by 19 species in the mock community (upper point) and 8 genera are represented by a single species.

The sourdough bread mock community was composed of 27 strains, 26 belonged to Ascomycota phylum while one was a Basidiomycota. At the genus level, 7, 5 and three species belonged to *Kazachstania, Pichia* and *Candida* spp. respectively, while all others belonged to a distinct genus. The choice of these species was based on a review paper on sourdough yeasts (Von Gastrow et al., 2023).

The cheese mock community contained 25 strains including 19, 5 and one from the Ascomycota, Mucoromycota and Basidiomycota phylum, respectively. Six strains belonged to *Penicillium*, 4 to *Mucor* and 2 to *Clavispora* spp. while the others belonged to a distinct genus. This choice of fungal species was based on the reviews by Montel et al. 2014 and Irlinger et al., 2015.

Among the 40 strains composing the fermented meat mock community, 36 and 4 belonged to the Ascomycota and Basidiomycota phylum, respectively. Nineteen, 4, 3, 2, 2 and 2 strains were affiliated to *Penicillium, Yarrowia, Cladosporium, Aspergillus, Rhododotorula* and *Candida*, respectively. All others belonged to a distinct genus. This choice was based on litterature data on fermented meats (Coton et al., 2021; Franciosa et al., 2021; Berni, 2014 and Selgas and Garcia, 2014).

For the wine mock community, it contained 60 strains belonging to Ascomycota (45 strains) and Basidiomycota (15 strains). Six strains were affiliated to *Hanseniaspora*, 6 to *Pichia*, 5 to *Candida*, 4 to *Rhodotorula*, 3 to *Papiliotrema* and 2 to *Clavispora, Cystobasidium, Metschnikowia* and *Meyerozyma*. All others belonged to a distinct genus. The selection was made from a review of papers investigating grape and wine microbiota (Setati et al., 2012; Rossouw and Bauer, 2016; Jolly et al., 2003; Setati et al., 2015; Bokulich et al., 2013 and Garofalo et al., 2016).

Twenty-seven of the 118 strains were common to at least 2 different mock communities (Bread, Cheese, Meat, Wine). One, i.e. *Torulaspora delbrueckii*, was present in all mocks.

### DNA extraction from single strains

#### Bread and wine

Each strain was grown overnight at 25°C in 15 mL of YEPD before centrifuging for 10 minutes at 1,500 × g. The cell pellet was resuspended in one mL of sterile water and transferred to a 2 mL tube. After a second centrifugation at 11,800 × g for 2 minutes, the pellet was resuspended in the yeast cell lysis solution from the MasterPure Yeast DNA extraction kit (Epicentre) and DNA was extracted according to the kit procedure.

#### Cheese

Each strain was grown overnight at 25°C under agitation at 200 rpm in 10 mL of YEGC. One mL of the culture was centrifuged at 10,000 × g for 10 minutes and the cell pellet was used for DNA extraction using the FastDNA SPIN Kit (MP Biomedicals).

#### Fermented meat

DNA was extracted from scraped colonies for yeasts or mycelial plugs for molds using the FastDNA SPIN Kit (MP Biomedicals) according to the manufacturer’s instructions.

### Mock community design

For each food environment (bread, wine, cheese, fermented meat), two different mock communities were prepared, a “DNA” mock community and a “PCR” mock community. For the DNA mock community, genomic DNA from each strain was quantified using the Qubit DNA Broad Range assay (ThermoFisher Scientific), diluted to the same concentration (10 ng/μL) and pooled. Then, the four markers were amplified in separate reactions from 20 ng of pooled DNA. For the PCR mock community, the four markers were individually amplified from each strain using 20 ng of genomic DNA as input, and PCR products were quantified using the Qubit DNA Broad Range assay (ThermoFisher Scientific). Then, 300 ng of PCR-product from each strain were pooled and diluted to a final concentration of 10 ng/μL before metabarcoding analysis. All mock community samples were prepared in triplicate.

### Real samples preparation

DNA extraction was performed for each food environment (bread, wine, cheese, fermented meat) according to different protocols adapted for each matrix. DNA concentration was determined using a Qubit fluorometer (Life Sciences) according to the Broad Range DNA assay kit protocol.

#### Bread

Three types of sourdough coming from different French bakeries were analyzed. All sourdoughs were made of wheat flour. Sourdough 1 was sampled from an artisanal bakery in Azillanet (Occitanie Region), sourdough 2 from a local baker in Assas (Occitanie region) and sourdough 3 from a local baker in Amilly (Centre-Val de Loire region). For each sourdough, DNA extraction was performed from 200 mg of three independent samples using the MO BIO’s Powersoil DNA isolation kit procedure (Qiagen 12888-100) as described previously (von Gastrow et al., 2022).

#### Cheese

Three ready-to-consume cheeses, namely Saint-Nectaire, Livarot and Epoisses, were analyzed. For each cheese type, three independent cheeses from the same producer were purchased on the same date. Rind was gently separated from the core using sterile knives, and only the rind fraction was analyzed. Rind samples were diluted 1:10 (w/v) in sterile distilled water and homogenized with an Ultra Turrax® (Labortechnik) at 8,000 rpm for 1 min to obtain a homogeneous mixture. DNA extraction was performed on 0.5 mL using the bead beating-based protocol detailed in a previous study (Dugat-Bony et al., 2015).

#### Fermented meat

DNA extractions were performed on casing samples obtained from French fermented sausages as described previously (Coton et al., 2021). Briefly, 5 cm × 1 cm casing samples were mixed with 9 mL sterile Tween (0.01% v/v) water followed by vigorous vortexing, before removing the casings. After centrifugation at 8,000 × g for 15 min, cell pellets were stored at −20°C until use. For DNA extractions, slightly thawed cell pellets were resuspended in 500 μL yeast lysis solution, divided in two, and DNA extracted using the FastDNA spin kit (MP Biomedicals) as described by the manufacturer. After extraction, DNA samples were purified using the DNeasy Tissue Kit silica-based columns (Qiagen) according to the manufacturer’s instructions.

#### Wine

Cells from 1 liter of a grape must were collected after centrifugation for 15 min at 10,000 × g. The pellet was resuspended in 10 mL YPD supplemented with 20% glycerol and stored at -80°C. Three samples of Sauvignon (1) and Viognier (2) grape must from the INRAE experimental wineries in Gruissan (France) were chosen. For DNA extraction, 1 mL of this cell suspension was sampled and centrifuged, then cells were resuspended in 1 mL of freshly prepared PBS supplemented with 1% Polyvinylpyrrolidone 25 to remove polyphenolic compounds that could further inhibit target amplification. After a second centrifugation at 15,000 × g for 10 minutes, DNA was extracted with the DNeasy Plant kit (QIAGEN, Hilden, Germany) with some modifications. The pellet was resuspended in 0.5 mL AP1 buffer and 4 μL of RNAse A solution and 300 μL of 0.3 mm glass beads were added. Cells were disrupted in a Precellys grinder (6,000 rpm, 320 seconds - 3 times). After centrifugation for 5 minutes at 15,000 × g, the supernatant was used for the downstream DNA extraction steps according to the manufacturer’s instructions. The resulting DNA samples were then used for metabarcoding.

### Library preparation and sequencing

Target markers were amplified with the primers presented in Table 2. The ITS1, ITS2, D1/D2 and RPB2 regions were amplified with the primers F (CTTTCCCTACACGACGCTCTTCCG-forward primer sequence) and R (GGAGTTCAGACGTGTGCTCTTCCG-reverse primer sequence) using 30 amplification cycles with an annealing temperature of 48 or 55°C (Table 2), 0.5 U MTP Taq (Sigma-Aldrich), 1.25 μL each primer (20 μM), 1 μL dNTP (10 μM each) in 50 μL final volume.

**Table 2.**
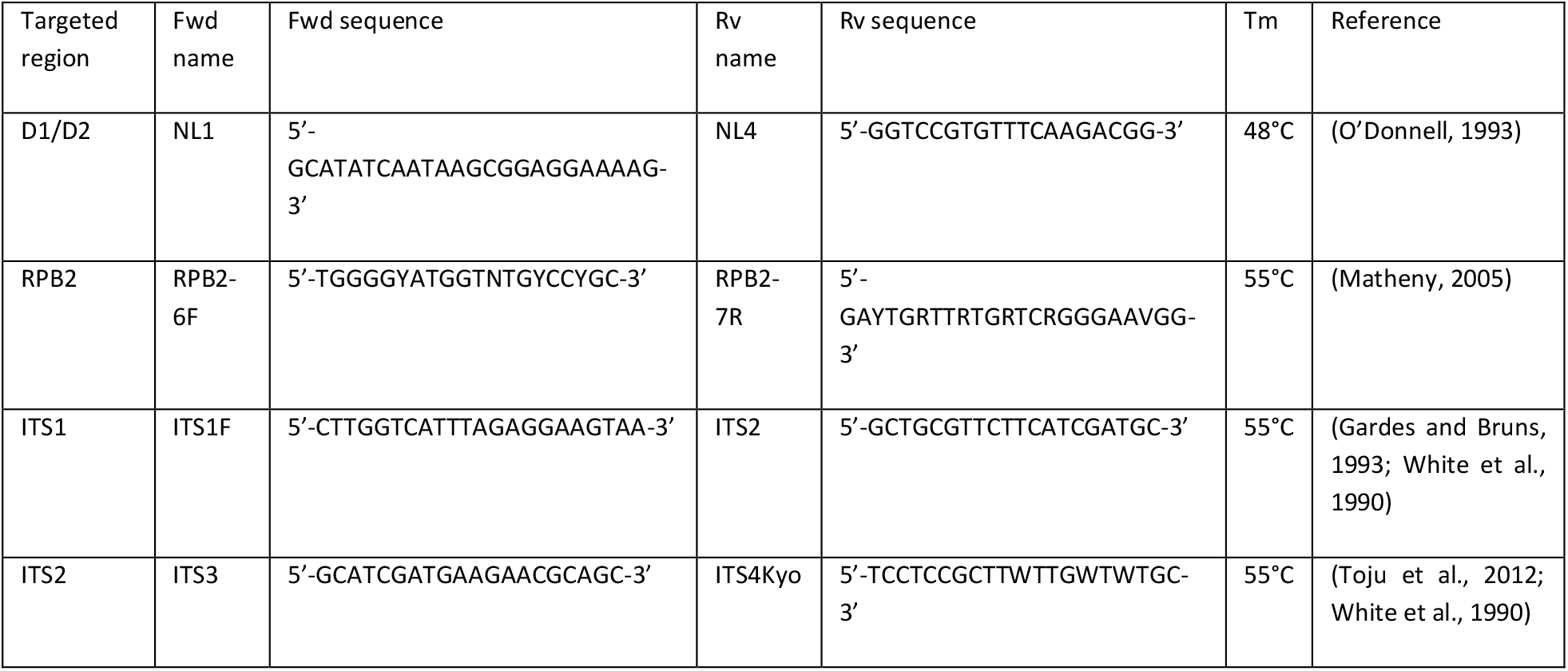
Description of the primers used to target ITS1, ITS2, D1/D2 and RPB2 regions.

Single multiplexing was performed using an in-house 6 bp index, which was added to the reverse primer during a second PCR with 12 cycles using forward primer (AATGATACGGCGACCACCGAGATCTACACTCTTTCCCTACACGAC) and reverse primer (CAAGCAGAAGACGGCATACGAGAT-index-GTGACTGGAGTTCAGACGTGT). The resulting PCR products were purified and loaded onto the Illumina MiSeq cartridge according to the manufacturer instructions, and paired-end read sequencing was performed for 2 × 250 cycles. The quality of the run was checked internally using PhiX as a control, and then each paired-end sequence was assigned to its sample with the help of the previously integrated index. The sequencing data from this study are available in NCBI SRA repository under the Bioproject number PRJNA685292.

### Bioinformatic analysis

#### Construction of the reference databank for mock community analysis

For each reference strain (118 in total), the sequence of the four phylogenetic markers (ITS1, ITS2, D1/D2, RPB2) was obtained from public databases or from unpublished sequences obtained in our labs. Only 3 RPB2 sequences were not available and missing due to PCR amplification failure (*Cryptococcus neoformans, Mucor lanceolatus* and *Rhodotorula glutinis*). The length distribution of the 469 sequences is represented in Figure 2. As expected, variations in length are visible, ranging from 70 to 835 bp. On average, ITS1 sequences are shortest, followed by ITS2, D1/D2 and RPB2 sequences.

**Figure 2.**
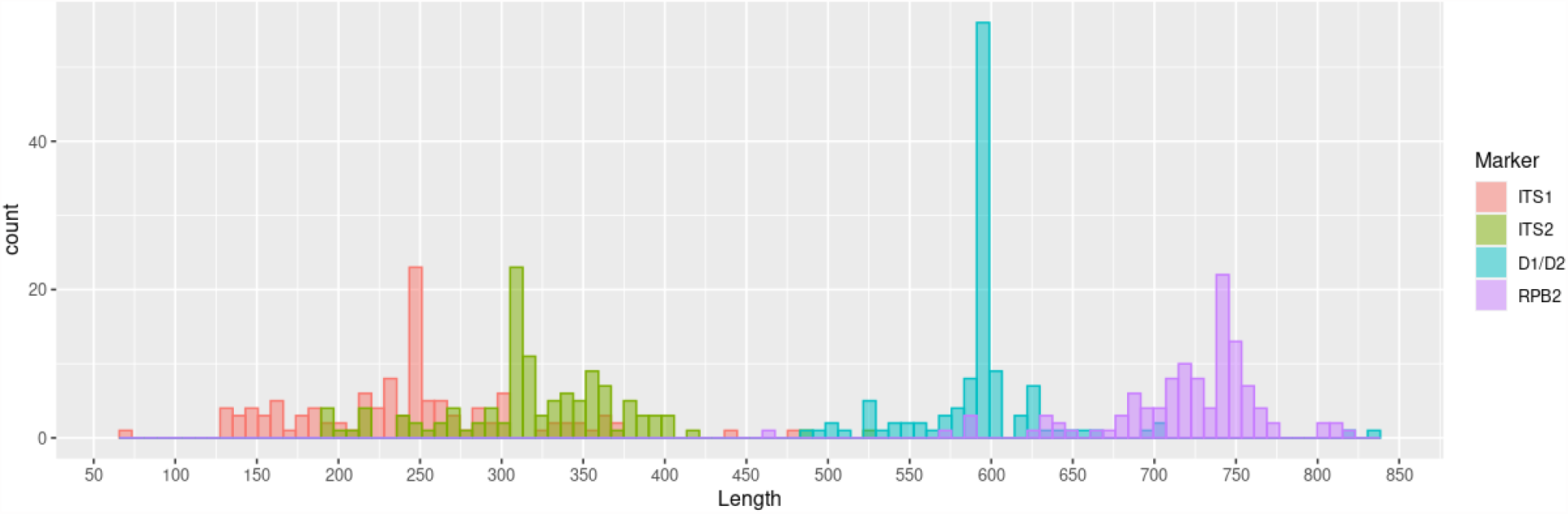
Sequence length distribution of the four markers for the 118 strains present in mock samples inferred from the reference database..

From all sequences and for each marker, we built phylogenetic trees with FastTree (Price et al., 2010) and Phangorn R package (Schliep, 2011) after a multiple alignment of sequences with Mafft (Katoh et al., 2009).

Figure 3 shows the diversity of the 118 ITS1 sequences from a phylogenetic point of view. Phylogenetic trees for other markers are available on Recherche Data Gouv platform (https://doi.org/10.57745/AZNJFE).

**Figure 3.**
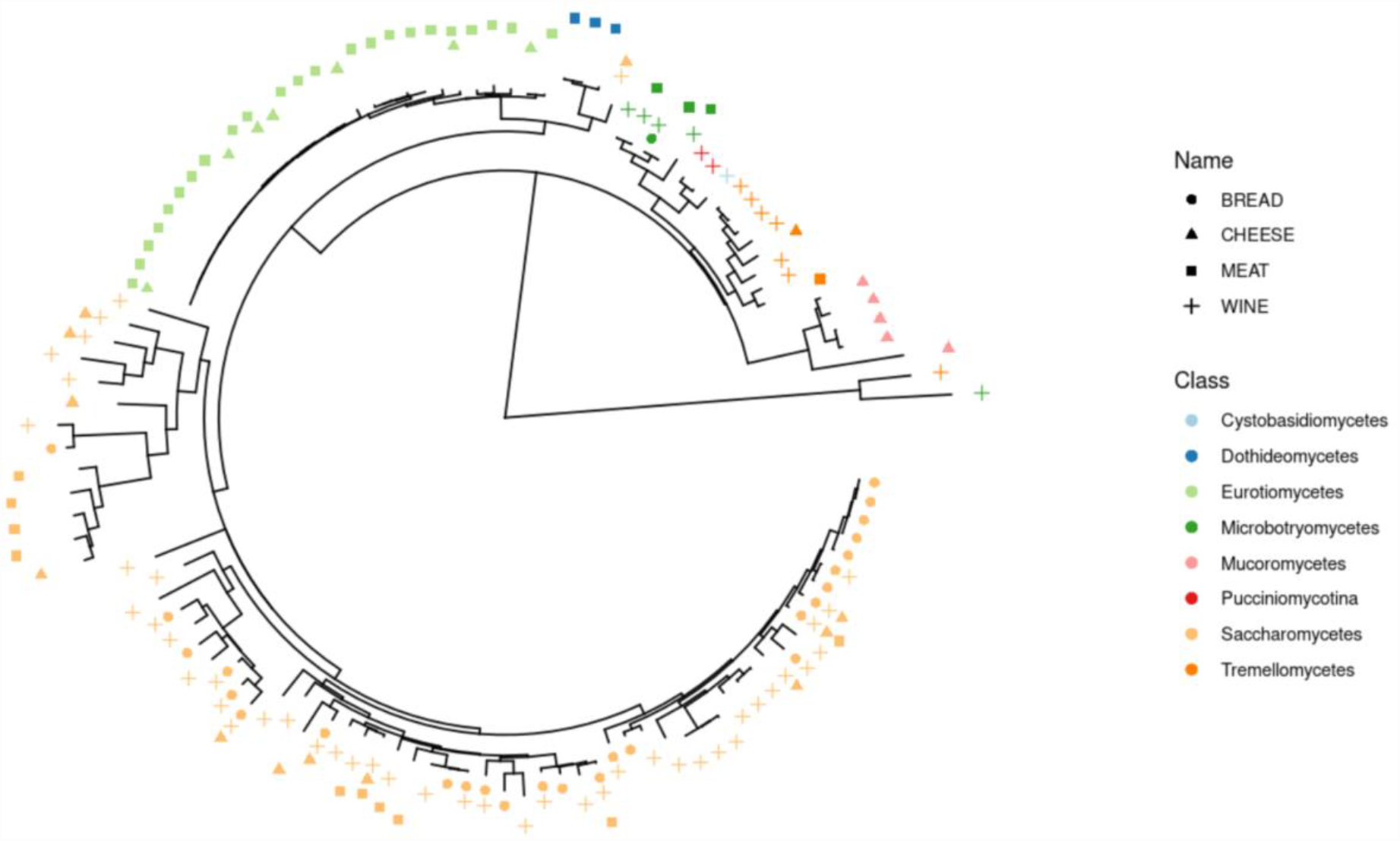
Phylogenetic tree built from D1/D2 sequences of the 118 strains of the mock samples, colored by Class. Shapes indicate the presence in bread, cheese, fermented meat or wine mock samples.

Database construction was crucial for benchmarking and was necessary in our case because many sequences are absent in public databanks. Figure 4 shows the best hits of the 469 sequences after a blast against each representative databank (UNITE v9.0 for ITS, SILVA v138 for D1/D2 and nt release 2021-07-30 for RPB2). If the sequence is present in the databank, the corresponding point is at the top right of the graph. If no hit is found, the corresponding point is at the bottom left of the graph. In the middle of the graph are points corresponding to sequences for which the percentage of identity and coverage are less than one hundred percent. This result illustrates the incompleteness of public databases for food ecosystems, particularly for RPB2 sequences.

**Figure 4.**
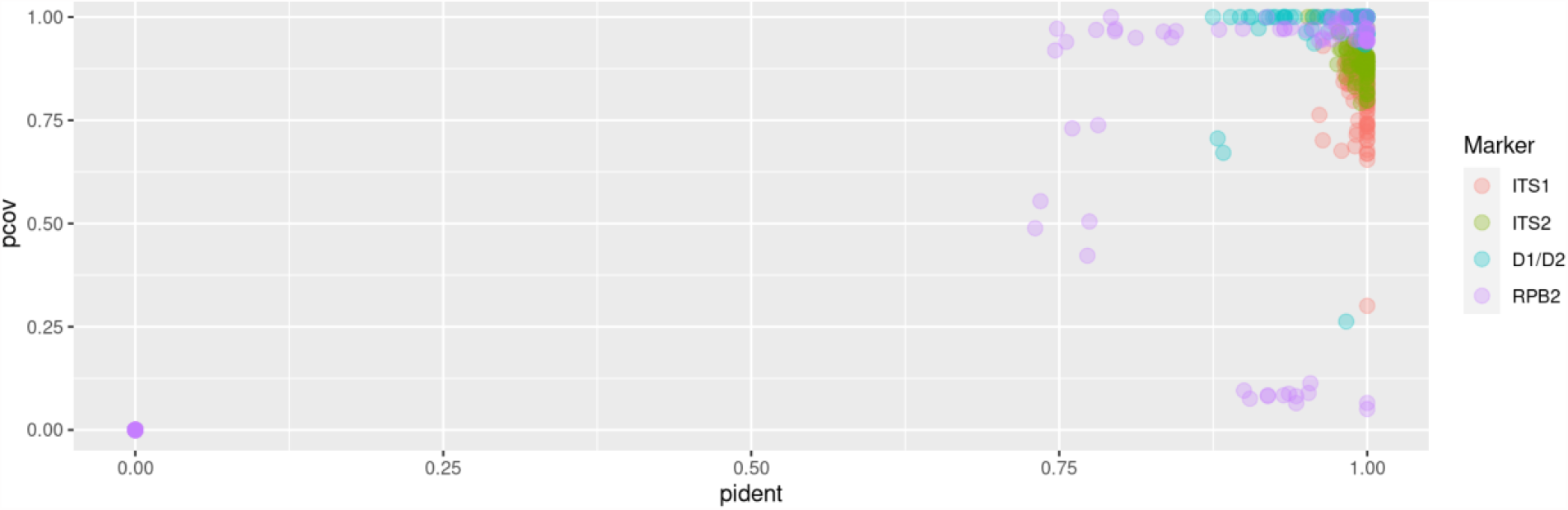
Representation of blast results (identity and coverage percentage) of the 468 sequences against dedicated databases (UNITE for ITS, SILVA for D1/D2 and nt for RPB2). The percentage of coverage and percentage identity are shown on the Y and X axis, respectively.

#### Benchmark of metabarcoding approaches

Full codes and figures are available on Recherche Data Gouv platform (https://doi.org/10.57745/109NNP). We used FROGS (Bernard et al., 2021), USEARCH (Edgar, 2010), QIIME2 (Bolyen et al., 2019) and DADA2 (Callahan et al., 2016) following their own guidelines, and a custom combination of DADA2 and FROGS that we named DADA2_FROGS.

One strength of FROGS is the ability to deal with overlapping and non-overlapping reads at the same time. This tool processes ITS1, ITS2, D1/D2 and RPB2 markers equally by a preprocess step (*preprocess*.*py*) to merge paired-end reads or create “artificial” sequences if they do not merge. In this case, R1 and R2 sequences are concatenated with a stretch of Ns in the middle. The subcommands *preprocess*.*py, clustering*.*py* with parameters *--fastidious* and *--distance 1, remove_chimera*.*py, otu_filters*.*py* with parameter *--min-abundance 0*.*00005, itsx*.*py, affiliation_OTU*.*py* and *affiliation_filters*.*py* with parameters *--min-blast-coverage 0*.*9, --min-blast-identity 0*.*9* and *--delete* were used.

DADA2 is a widely used tool for metabarcoding analyses. It infers exact amplicon sequence variants (ASVs) from amplicon data, resolving biological differences of even 1 or 2 nucleotides. Cutadapt and then the functions *filterAndTrim (maxN = 0, maxEE = 2, truncQ = 2, minLen = 50, rm*.*phix = TRUE), dada and mergePairs* were used. By default, DADA2 does not deal with overlapping and non-overlapping reads at the same time. We used single-end and paired-end modes (DADA2-se and DADA2-pe, respectively). For DADA2-se, only R1 reads were taken into account and only overlapping reads for DADA2-pe. Then, for both strategies, *makeSequenceTable, removeBimeraDenovo* and *assignTaxonomy* functions were finally used.

In the same way, QIIME2 was used in single and paired-end modes (QIIME-pe and QIIME-se). The commands used were *qiime cutadapt, qiime dereplicate-sequences*and then we performed an open-reference clustering using the *qiime vsearch cluster-features-open-reference* command to build OTUs with the parameter *--p-perc-identity 0*.*99*. Chimera were removed with *vsearch uchime-denovo* command and the taxonomic affiliation was done with *qiime feature-classifier classify-sklearn*.

For USEARCH, we followed the instructions provided by the author on his website (https://www.drive5.com/usearch/manual/global_trimming_its.html) by taking into account merged sequences and 5′ R1 reads of non-overlapping reads and used successively the parameters *-fastq_mergepairs,-search_oligodb, -fastq_filter, -fastx_uniques, -unoise3 and -otutab* to produce ZOTUs and *-sintax* for taxonomic affiliation.

For the DADA2_FROGS strategy, the DADA2 recommendations were followed until obtaining the ASV table (cutadapt, *filterAndTrim, dada, mergePairs* and *makeSequenceTable* functions). At this step, we followed the FROGS guidelines after the clustering step: remove chimera (*remove_chimera*.*py*), ITSx (itsx.py) for ITS data and taxonomic affiliation (affiliation_OTU.py). The aim was to benefit from the denoising algorithm that is, in theory, able to produce high-resolutive ASVs. As we wanted to keep merged and unmerged reads, we kept them by using the returnRejects parameter of the dada2 mergePairs function.

For each tool, we used our internal database available on Recherche Data Gouv platform (https://doi.org/10.57745/AZNJFE), consisting of our mock sequences, for taxonomic affiliation of the OTUs/ASVs/ZOTUs, as described above. No additional sequence was added to avoid unnecessary noise to analyze the different mock communities.

Different metrics were computed in order to compare the above-mentioned methods: (i) the divergence rate, computed as the Bray-Curtis distance between expected and observed abundance profiles at the species level; (ii) the number of false-negative taxa (FN) corresponding to the number of expected taxa that were not recovered by the method, (iii) the number of false positive taxa (FP) corresponding to the number of recovered taxa that were not expected, (iv) the number of true positive taxa (TP) corresponding to the number of recovered taxa that were expected. From these metrics we computed the precision (TP/(TP+FP)) and the recall rate (TP/(TP+FN)). Finally, as we knew the exact expected sequences, we computed the number of sequences perfectly identified (OTUs/ASVs/ZOTUs with nucleic sequence was strictly identical to the known reference sequence). For long sequences (i.e. > 500 bp), the middle was not sequenced and only the sequenced part was used for taxonomic affiliation. In this case, 100% identity between the reference and the OTU/ASV/ZOTU resulted in a perfect identification.

#### Analysis of real samples

DADA2_FROGS, the bioinformatics approach with the best results from mock samples, was used to analyze real samples (Code and figures are available on Recherche Data Gouv platform, https://doi.org/10.57745/ENE09G). For the taxonomic affiliation of these samples (composition unknown), and for each marker, we added the 118 sequences from our mock communities to Unite (v. 9,0) (Rolf Henrik Nilsson et al., 2019) for ITS data and SILVA (v. 138) (Quast et al., 2013) 28S rDNA sequences for D1/D2. For RPB2, we needed to build an in-house database because no dedicated one was publicly available to the best of our knowledge. We first extracted sequences from the “Fungi” division from NCBI nt databank (release 2021-07-30) (Sayers et al., 2022) using taxonkit (v. 0.6.0) (Shen and Ren, 2021) and then used cutadapt (Martin, 2011) with RPB2 primers to target sequences of interest. The databases were composed of 206,184 ITS sequences, 16,293 D1/D2 sequences and 13,055 RPB2 sequences.

For each ASV obtained, the taxonomic affiliation was manually checked and corrected when needed. More precisely, ASVs were blasted against different databases (e.g., NCBI, YEASTIP) to confirm or correct the affiliation, and we removed some ASVs (remaining chimera, contaminations). When taxonomic resolution at the species level was not possible (identical sequences between two or several species), we defined groups of species and labeled ASVs accordingly.

This manual curation step was performed for the most abundant ASVs (an abundance of at least 150 by marker and food ecosystem).

## Results

In this study, we compare the efficiency of 4 barcodes (ITS1, ITS2, D1D2, RPB2) and seven bioinformatic workflows to detect the species in microbial community of 4 fermented products (bread, wine, cheese, fermented meat) using mocks and real samples. The phylogenetic diversity of fungal species analyzed is illustrated Figure 3.

### Choice of the most accurate bioinformatic approach

Our benchmark of tools was only performed on mock community samples, and the results of the four markers were analyzed together (Figure 5).

**Figure 5.**
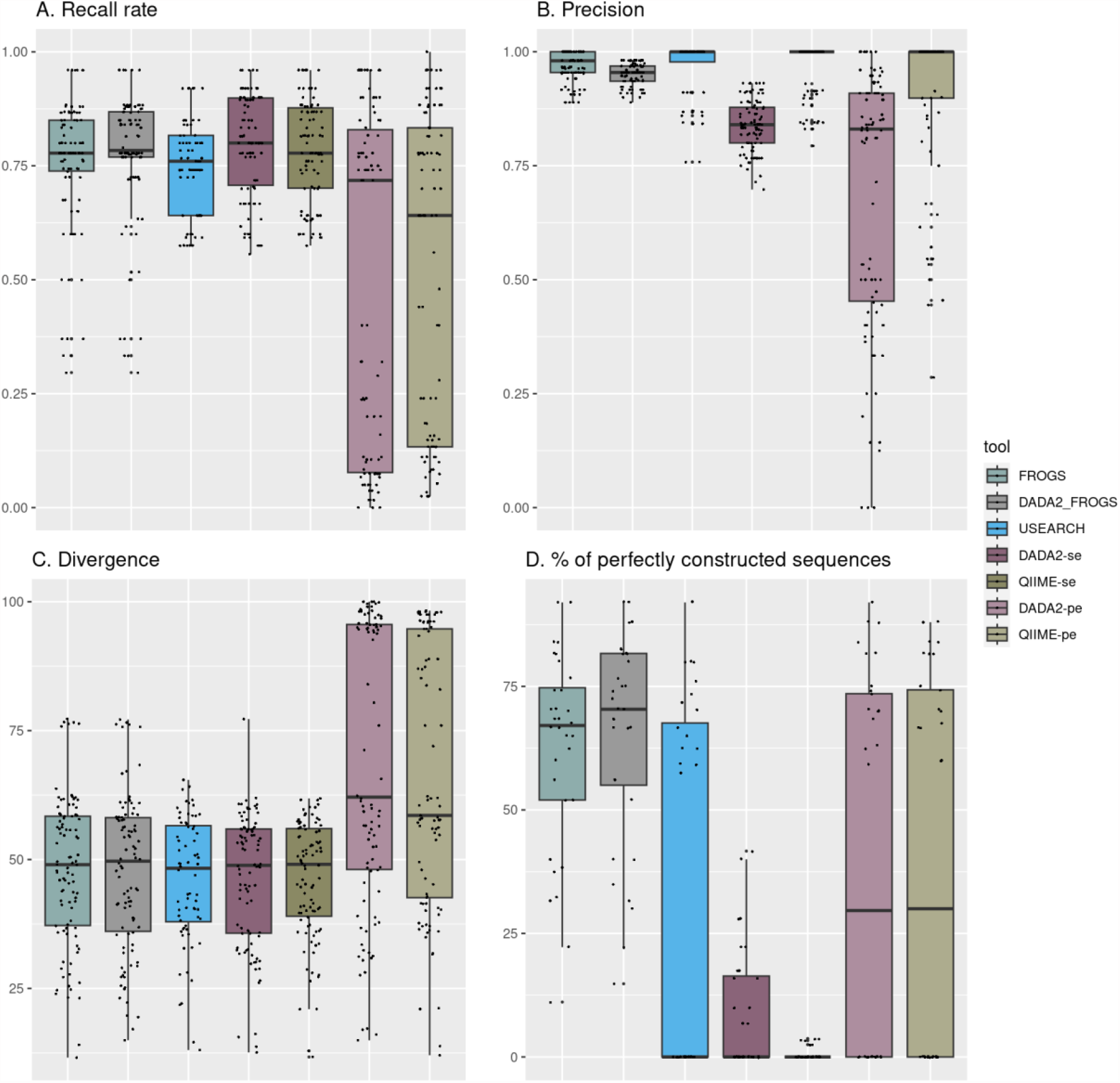
Quality parameters obtained with the seven bioinformatics pipelines. A) Recall rate (TP/(TP+FN)) reflects the capacity of the tools to detect expected species. B) Precision (TP/(TP+FP)) shows the fraction of relevant species among the retrieved species. C) Divergence rate is the Bray-Curtis distance between expected and observed species abundance. D. Percentage of perfectly reconstructed sequences is the fraction of predicted sequences with 100% of identity with the expected ones.

### Recall rate

The median recall rate (sensitivity), reflecting the capacity of tools to detect expected species, is between 0.75 and 0.8 for FROGS, DADA2_FROGS, USEARCH, QIIME-se and DADA2-se. It is lower (∼0.5) for QIIME-pe and DADA2-pe, due to the fact that these methods always reject reads if they do not overlap (all D1/D2 and RPB2 sequences and some ITS1/2 were always missing).

### Precision

Regarding precision, the four methods yield values of 0.95-0.97 (FROGS, DADA2_FROGS, USEARCH and QIIME-se). QIIME-pe is slightly less efficient (0.92), and both DADA2-se (0.84) and DADA2-pe (0.69) are worse.

### Divergence rate

The divergence rate is computed as the Bray-Curtis distance between expected and observed abundance profiles at the species level. It therefore reflects the ability of the tool to detect species in the right proportions. The divergence rates obtained in this study are very high and, as expected, are lower for PCR mocks, as we can see in Figure 7. FROGS, DADA2_FROGS, USEARCH, QIIME-se and DADA2-se yield equivalent results for this indicator (∼46-47% on average) while DADA2-pe and QIIME-pe show higher divergence rates (65-69%).

### Reconstruction of sequences

DADA2_FROGS is able to reconstruct more sequences than the other methods. Indeed, 79.2% of expected sequences are found without errors. FROGS is very close with 75.9%, followed by DADA2-pe (75.1%) and QIIME-pe (74.9%). USEARCH (66%), DADA2-se (17.6%) and QIIME-se (1.2%).

Overall, the results obtained with the four indicators reveal that the DADA2_FROGS approach performs the best for analyzing ITS1, ITS2, D1/D2 and RPB2 mock samples. We thus selected this approach for all subsequent analysis. Nevertheless, it should be noted that the FROGS tool also performs well as it yields indicator values that are similar to DADA2_FROGS. The main difference is due to species harboring very similar sequences, such as those belonging to Penicillium spp.

### Effect of amplicon length on the detected relative abundance

The ITS1 and ITS2 amplicon size is highly variable depending on the considered fungal species, as observed for those included in our mock communities (Figure 2). The effect of amplicon size on the relative abundance of the different species was evaluated using the PCR mock dataset (Figure 6) (code and figures are available on Recherche Data Gouv platform: https://doi.org/10.57745/APNOH8).

**Figure 6.**
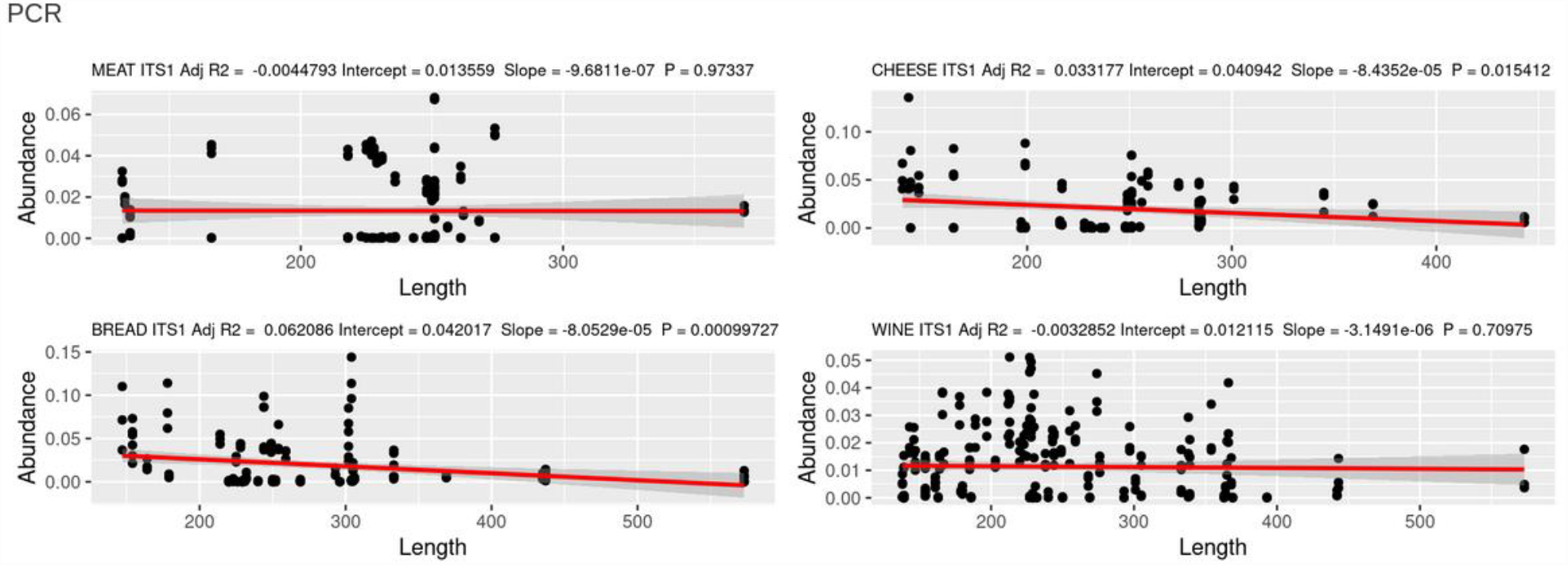
Effect of ITS1 amplicon size on the relative abundance of the It is rather likely that all primers have missmatches with certain groups of fungi.detected ASVs in the PCR mock datasets.

**Figure 7.**
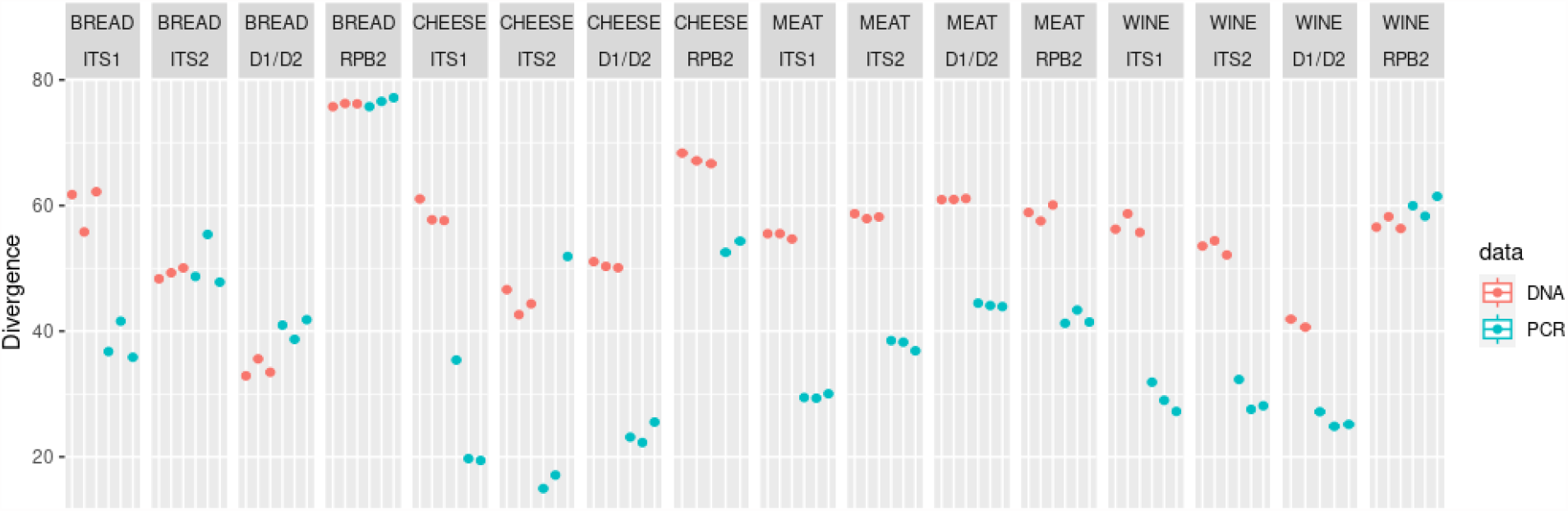
Divergence at species level for each tested barcode marker and mock community. DNA mock communities are colored in red and PCR mock communities are colored in blue.

A significant negative relationship is observed between amplicon length and relative abundance in two out of four tested mock communities for ITS1 (cheese and bread, but not meat and wine) and ITS2 (meat and wine, but not cheese and bread). Furthermore, the determination coefficient (R2), which indicates the proportion of variation in the relative abundance data that is predictable from the amplicon length, is comprised between 0.013 and 0.085. This parameter therefore only has a limited impact on the observed proportions when using ITS1 and ITS2 as barcode markers.

### Comparison of markers

To compare the capacity of each marker to correctly reflect the fungal diversity present in fermented food samples, we only focused on the results obtained with the DADA2_FROGS approach. Code and figures are available on Recherche Data Gouv platform (https://doi.org/10.57745/X6AXA6). Figure 7 shows the divergence rate obtained for each marker and mock community type (DNA or PCR mock communities).

Figure 8 shows the amount of absent, partially and perfectly reconstructed sequences for each ecosystem.

**Figure 8.**
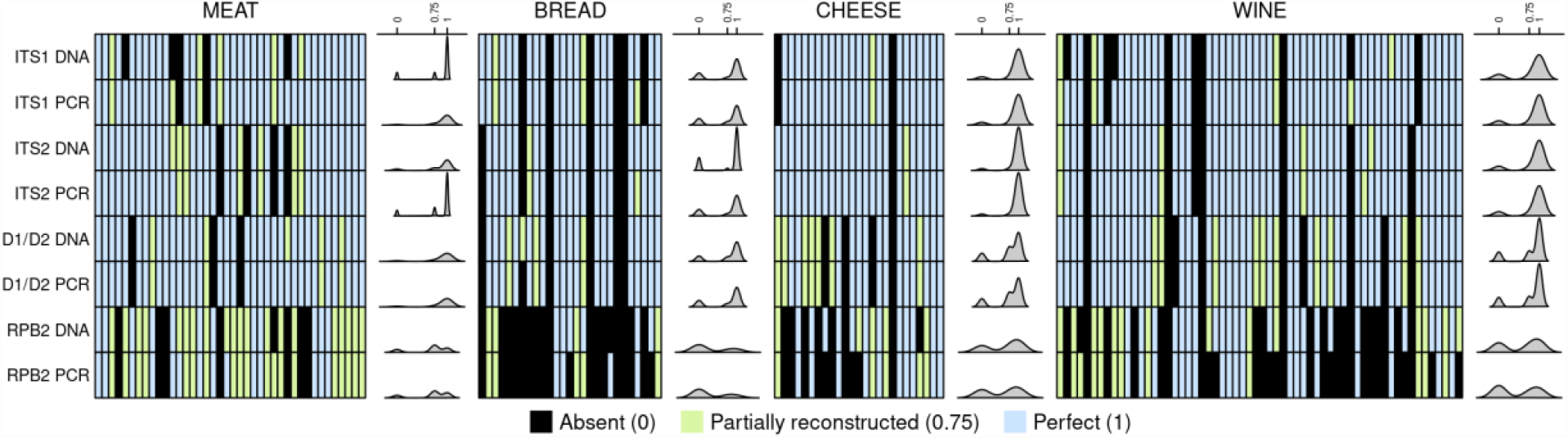
Heatmap of expected species for each mock community and barcode marker using the DADA2_FROGS approach. Dark green represents perfectly reconstructed sequences (score of 1), light green partially reconstructed sequences (score of 0.75) and orange species that are missing (0). Density of results is indicated for each line, representing the ability of markers to be efficient.

#### Bread

Regarding the sourdough bread mock community, divergence varies according to the tested barcode marker. While RPB2 presents the highest divergence (75%), D1/D2 shows the lowest (34%-42%) while ITS1 and ITS2 show an intermediate level of divergence. Contrary to what was found for cheese and meat mock communities, divergence is not higher for the mock community DNA mixture than for the mock community PCR mixture, except for ITS1 where the PCR mock mixture presents 35% divergence while the DNA mock mixture ranges from 55% to 61%.

ITS1 and ITS2 perform equally well in terms of false negatives (n=6), false positives (n=1) and true positives (n=21). D1/D2 exhibits one less false negatives (n=5), one more true positives (n=22) and one more false positives than both ITS barcode markers. RPB2 is the worst performing marker with the highest number of false negatives (n=12-14), and the lowest number of true positives (n=8-10).

These results show that D1/D2, ITS1 and ITS2 are all relevant for the analysis of sourdough microbiota but it cannot be concluded which one is best.

#### Cheese

Regarding the cheese mock community, D1/D2, ITS1 and ITS2 show comparable performance in terms of divergence and are more accurate than RPB2. It is noteworthy that divergence is higher for DNA (between 45 and 70%) than PCR (between 15 to 50%) mock communities. ITS2 exhibits less false negatives (only one) and more true positives (24/25) than other markers. However, it generates two false positives with the DNA mock community and one with the PCR mock community samples while D1/D2 and RPB2 generates one false positive with both DNA and PCR samples. Altogether, these results indicate that, according to the four markers used in this study, ITS2 provides the best compromise to accurately profile cheese fungal communities.

#### Fermented meat

Regarding the fermented meat mock community, all tested markers show comparable performance in terms of divergence, ranging from 52 to 56% and 23 to 35% for the mock DNA and mock PCR mixture, respectively. As observed for the cheese mock community, divergence is 1.5-2 times higher for the mock DNA mixture than for the mock PCR mixture, the lowest divergence being observed with ITS1 marker in the mock PCR mixture (23% divergence). D1/D2 exhibits the highest number of false negatives (n=8) as compared to other markers while for PCR mock community mixture, ITS1 and RPB2 exhibit 4 and 5 false negatives, respectively. Concerning the true positive metric, all markers perform equally well with between 32 and 33 true positives out of 40 expected species for both mock DNA and PCR mixtures with the exception of ITS1 marker in the mock PCR mixture which yields 36 true positives. Based on the above mentioned results, we conclude that the ITS1 barcode is slightly more accurate for profiling fermented meat fungal communities, although ITS2 and RPB2 also performed well.

#### Wine

RPB2 marker does not detect most species (22/60 not found). In contrast, D1/D2, ITS1 and ITS2 display similar results to describe the mock community although not completely; ITS2 is slightly better than the other markers. For the latter three markers, similar performance in terms of divergence is obtained, but better for the DNA mock community than the PCR mock community. However, at least 7 species out of 60 are not identified. ITS2 is also shown to be the most efficient.

Similar to the cheese ecosystem, the ITS2 barcode was the most accurate to explore wine mycobiota, followed by ITS1.

### Analysis of real samples

We then compared the efficiency of the four barcodes to detect species in real samples in order to validate our mock results and take into account the fermented food matrix. Code and figures are available on Recherche Data Gouv platform (https://doi.org/10.57745/ENE09G).

#### Bread

Metabarcoding results from the wheat sourdough sample analyses showed that hits with identities above 80% were not detected using the RPB2 marker. Besides fungal DNA, the other three markers amplified plant DNA. The number of plant DNA hits was much higher for ITS2 and D1/D2 than ITS1. Moreover, ITS2 and D1/D2 markers amplified DNA from *Triticum* species (*Triticum aestivum, Triticum monococcum, Triticum durum*), and crop weeds, such as *Viciae* sp., *Gallium* sp. and *Calystegia* sp., often found in cereal fields. The ITSx tool automatically removed plant-derived ASVs in the final ITS1 and ITS2 ASV table whereas those from the D1D2 dataset had to be manually removed.

Regarding filamentous fungi, several genera were not detected with D1/D2 including *Aspergillus, Aureobasidium* and *Tilletia*. Regarding mycotoxin-producing wheat pathogens (e.g., *Fusarium* and *Penicillium* spp.) or species involved in negatively impacting grain and flour quality for bread making (e.g., rotten fish smell due to *Tilletia* sp.), results showed that the ITS1 barcode did not detect *Penicillium* spp. contrary to D1/D2 and ITS2 barcode markers. On the other hand, ITS1 allowed a better resolution to the species level within the *Fusarium* and *Tilletia* genus.

Regarding fermenting yeast, the well-known bakery yeast *Saccharomyces cerevisiae* was detected by all three markers. In contrast, *Wickerhamomyces anomalus*, frequently encountered among dominant yeast species in sourdoughs worldwide, was only detected by the ITS1 and ITS2 markers although not consistently. It was found in all sourdough samples using the ITS1 marker but only in two out of the three sourdough samples using the ITS2 marker.

Based on the comparison of the RPB2, ITS1, ITS2, D1/D2 markers on real sourdough samples, ITS1 is the best adapted marker to describe sourdough mycobiota as lower reads due to plant DNA were observed and the best detection of sourdough fungal species was obtained.

#### Cheese

At the genus level, D1/D2, ITS1 and ITS2 all detected *Geotrichum* and *Debaryomyces* as the major fungal genera present on the surface of the three studied cheeses, contrary to RPB2, which placed *Debaryomyces* and *Kluyveromyces* as the dominant taxa and exhibited high variations between biological replicates. So, we decided to exclude RBP2 from the comparison. Regarding the three other markers, some important discrepancies were observed. First, *Yarrowia* and *Mucor* species were only detected in real cheese samples with D1/D2 and ITS2. These species are among the most prevalent fungi in cheese products. Secondly, within *Geotrichum*, ITS1 detected ASVs affiliated with *G. candidum* but also with *Geotrichum* sp. while both D1/D2 and ITS2 only detected *G. candidum* species. Thirdly, within the *Mucor* genus, better species level resolution was observed with ITS2 compared to D1/D2. Similarly, it was possible to correctly assign the species *Kluyveromyces lactis* to the corresponding ASV when using ITS1 and ITS2, but the one obtained with D1/D2 could not be differentiated between *K. lactis* and *K. marxianus*. As these two species are frequently used as ripening cultures in cheese production, it may be important to choose the ITS1 or ITS2 primers to discriminate between them. Finally, *Candida*, which aggregates with species from different genera including species with uncertain affiliations, was abundant in sample 1 (Saint-Nectaire cheese) based on both ITS regions but was only slightly detected with D1/D2. Altogether, these results indicate that ITS2 performs best to describe cheese fungal communities, followed by D1/D2.

#### Fermented meat

At the genus level, *Debaryomyces* and *Penicillium*, which are major genera on the casings of fermented meat, were detected with all tested barcode markers. Interestingly, *Kurtzmaniella* sp. (*“Candida” zeylanoides*) was only identified using D1D2 and ITS2 barcodes while *Yarrowia* and *Scopulariopsis* sp. were only identified using D1D2 and ITS2, and, D1D2, ITS2 and RPB2, respectively. Noteworthy, both genera were found in a much larger number of samples using the ITS2 barcode. At the species level, *D. hansenii* and *Penicillium nalgiovense* were found, as expected, to be the most dominant taxa by all 4 barcode markers. Noteworthy, an unambiguous assignation of ITS1 and ITS2 ASVs to *P. nalgiovense* was not possible as these ASVs shared 100% similarity with other related *Penicillium* species and with *Penicillium melanoconidium* for ITS1 and ITS2 ASVs, respectively. Among other major species in fermented meat, *P. salamii*, was only identified with ITS1, ITS2 and RPB2 barcodes while *P. nordicum*, a mycotoxin-producing fungus, was only found using RPB2. ITS2 was the only barcode that accurately identified *Yarrowia* species (i.e., *Y. deformans* and *Y. lipolytica*). Overall, among the different tested barcodes, ITS2 was the most efficient for identifying the majority of fungal meat species, the only exception being for the very diversified *Penicillium* genus.

#### Wine

In comparison to the other studied ecosystems, species diversity in grape must samples was much more complex. More than one hundred yeast and filamentous fungal species were detected including 33 that were part of the wine mock.

When comparing the main species encountered, striking differences were observed between ITS1, ITS2 and D1D2 results. *Aspergillus* and *Aureobasidium* were detected in all samples, although with large differences according to barcode. Nearly 50% was identified in the triplicates of sample 1 using ITS2 while only 25% with ITS1 and about 40% with D1D2 (higher abundances of *Aspergillus* than *Aureobasidium* were also noted). An opposite situation was observed for *Botrytis*. Indeed, this genus was much more abundant using ITS1 versus ITS2 and D1D2. In agreement with mock analysis results, *Starmerella bacillaris*, a major component of grape must microbiota, was detected at very low abundance in the different samples with ITS1, whereas this species was detected in all samples with ITS2 and D1D2, and represented up to 40% in sample 2 according to ITS2 read counts. In a similar manner, *Hanseniaspora uvarum* was poorly detected with ITS1, well detected with ITS2 and most abundant with D1D2. In contrast, *Metschnikowia* species were detected in a similar manner with ITS1 and ITS2 (more than 40% abundance in samples 3.1 and 3.2) despite the short length and the polymorphism of the amplified sequences.

In conclusion, the ITS2 barcode provides the most comprehensive description of grape must mycobiota.

## Discussion

The aim of this study was to compare bioinformatic tools and barcodes used to describe fungal communities in different fermented foods. Choosing the most robust barcode marker for accurate description of fungal communities is crucial. However, various challenges/biases have to be addressed or taken into account for their accurate characterization such as incompleteness of reference databases, low taxonomic depth and PCR amplification biases. In addition, dedicated pipelines also need to be evaluated. In the present study, we built mock fungal communities that gathered the most representative species of fermented meat, cheese, sourdough bread and grape must (wine). These mock communities were used to compare the performances of the main bioinformatic tools available to the scientific community (FROGS, USEARCH, QIIME and DADA2) as well as a combination of DADA2 and FROGS using reads obtained from four commonly used barcodes for fungal community assessment, i.e. ITS1, ITS2, D1D2 of the rDNA as well as RPB2. In addition, to compare these bioinformatic pipelines, we built an in-house database of barcode sequences as many sequences from major fungal species found in these fermented foods were missing in currently available databases. Finally, after selecting the best bioinformatic pipeline, we compared the performances of these four barcodes using real fermented food samples.

In the first part of this study, we compared several commonly used bioinformatic tools. By combining the denoising step of DADA2 followed by the FROGS pipeline, we defined a “universal” pipeline for all barcodes. It combines the advantages of FROGS (dealing with all amplicon lengths) and those of denoising approaches (best resolution and stable ASVs to compare datasets from different studies). Our pipeline avoids the pitfall of other tools in which targeting short barcodes rather than long ones is required (Brandon-Mong et al., 2015; Leray et al., 2013).

One of the main challenges in fermented foods is to characterize microbial communities at the species level including fungi. Food fungal communities are less diversified at the genus level but diversity within genera needs to be determined as it can be relatively high. Robust barcode markers are thus required to reach species level descriptions among genera. Protein-coding genes can be useful to reach this goal and among them, RPB2 is one of the most commonly used taxonomic marker. However, our results clearly highlighted the poor performance of this gene as a barcode, due to a lack of amplification of the barcode. This might be related to the choice of primers, which were originally designed for Basidiomycota (Matheny, 2005). To overcome this limitation, it would be worth designing new consensus primers suitable for Basidiomycota, Ascomycota and Mucoromycotina. Alternative protein-coding genes also need to be tested.

Besides protein-coding genes, rDNA barcodes provide a good global view of mycobiota. Previous studies compared ITS1 versus ITS2 (Bokulich and Mills, 2013) or ITS versus nuclear ribosomal large subunit (LSU) barcodes (Brown et al., 2014). All studies converged on the proposition of using ITS as the primary fungal barcode (Schoch et al., 2012). The LSU appeared to have superior species resolution in some taxonomic groups (Mota-Gutierrez et al., 2019), such as the early diverging lineages and ascomycete yeasts, but was otherwise slightly inferior to ITS (Schoch et al., 2012). ITS1 and ITS2 are, in general, more resolutive markers than D1D2, in particular for filamentous fungi (Mota-Gutierrez et al., 2019). ITS1 locus generally has the shortest mean amplicon lengths for all phyla, the smallest difference between Ascomycota and Basidiomycota amplicon lengths, and the highest species- and genus-level classification accuracy for short amplicon reads, arguing for the primacy of this locus, compared to ITS2 (Bokulich and Mills, 2013). However, in the present study, we found that none of the rDNA markers allowed us to unequivocally discriminate between all species in real fermented foods. For example, this was emphasized for several mold species, especially species belonging to *Penicillium* or *Pichia spp*.

While D1D2 is among the reference sequences for fungal taxonomy, we found that it had less discriminating power to differentiate species as compared to ITS1 and ITS2 barcodes. This agrees with the previous analysis of 9,000 yeast strains, showing that 6 and 9.5% of the yeast species could not be distinguished by ITS and LSU, respectively (Vu et al., 2016). Indeed, LSU is more conserved than ITS.

The ITS1 and ITS2 markers performed better than D1D2 but their performance varied according to the tested fermented foods. Indeed, while ITS2 performed better for cheese, meat and wine, ITS1 seems better for bread. Concerning the latter, ITS2 primers amplified the ITS from wheat and several weeds which hampered its efficiency for cereal-based products. Besides the fermented product being studied, the choice of the ITS barcode also depends on the expected species diversity in the ecosystem and may be driven by fungal species of interest. Moreover, choice of ITS primers can also be adapted to targeted species. Indeed, some ITS barcoding primers may have mismatches with the sequence of fungal species of interest, such as *Yarrowia* species (Ihrmark et al. 2012, Tedersoo and Lindahl, 2016). Finally, although ITS1 and ITS2 seem to be the best barcodes for distinguishing between species and, according to our results, their variation in size does not appear to introduce a large bias, their difference in size may hinder sequence alignment and therefore beta diversity estimates that take phylogenetic distances into account.

One of the limits addressed in the present study was the availability of a complete database adapted to the chosen fermented foods. We thus developed an in-house sequence database for all four major fermented foods in order to fill in this gap. This database significantly improved species level affiliations although manual curation was still required for some genera with complex taxonomy such as *Penicillium* spp. These results also illustrate the need to expand public databases with specific databases.

Metabarcoding is known to be a semi-quantitative method. It is considered to suffer from amplification biases caused by fragment length polymorphism. We did not find evidence for any correlation between amplicon size and read abundance. The divergence between the expected and observed frequency likely results from differences in copy number of rDNA genes (Sternes et al., 2017). Normalization of relative abundance by qPCR targeting a standard reference (Zemb et al., 2020) or by digital PCR (Floren et al., 2015; Zimmer-Faust et al., 2021) might also correct for DNA extraction bias.

In conclusion, although ITS2 appears as the most accurate barcode marker for fermented meat, cheese and wine samples and ITS1 for sourdough bread, no generic recommendation for all fermented food types can be made. This is mainly due to the fact that taxonomic resolution within some genera is not efficient which highlights the need to combine metabarcoding with culture-dependent analysis such as culturomic approaches. The availability of long-read technologies, like Oxford Nanopore Technologies or PacBio technology, provides the opportunity to sequence longer fragments of the fungal ribosomal operon, up to 6 Kb (18S-ITS1-5.8S-ITS2-28S) and to improve the taxonomy assignment of the communities up to species level (D’ Andreano et al., 2021) but their current cost is still a brake to replace short reads technologies. Shotgun metagenomics sequencing is also an alternative or a complementary method to study food fermentations (Leech et al., 2020). It may provide a less biased vision of food microbiota than metabarcoding (Sternes et al., 2017), a more comprehensive insight into the microbial composition, and functional potential but at a much higher cost for low abundant species.

## Acknowledgements

We are grateful to the INRAE MIGALE bioinformatics facility (MIGALE, INRAE, 2020. Migale bioinformatics Facility, doi: 10.15454/1.5572390655343293E12) for providing computing and storage resources. We thank Julien Lebrat, Stéphane Guezenec, Anne-Sophie Sarthou for technical assistance and Claire Vincent for her participation in building the reference databank. We also thank Mary and Loulou Bonneau, Stéphane Marrou, Anna and Maximilien for providing security for their sourdough.

## Data, scripts, code, and supplementary information availability

Sequencing data are available in NCBI SRA repository under the Bioproject number PRJNA685292. Scripts and code are available online on Gitlab (https://forgemia.inra.fr/migale/metabarfood).

Supplementary information is available online: https://doi.org/10.57745/AZNJFE, https://doi.org/10.57745/109NNP, https://doi.org/10.57745/ENE09G, https://doi.org/10.57745/X6AXA6 and https://doi.org/10.57745/APNOH8.

